## Supplementary Information for "Allosteric activation of cell wall synthesis during bacterial growth"

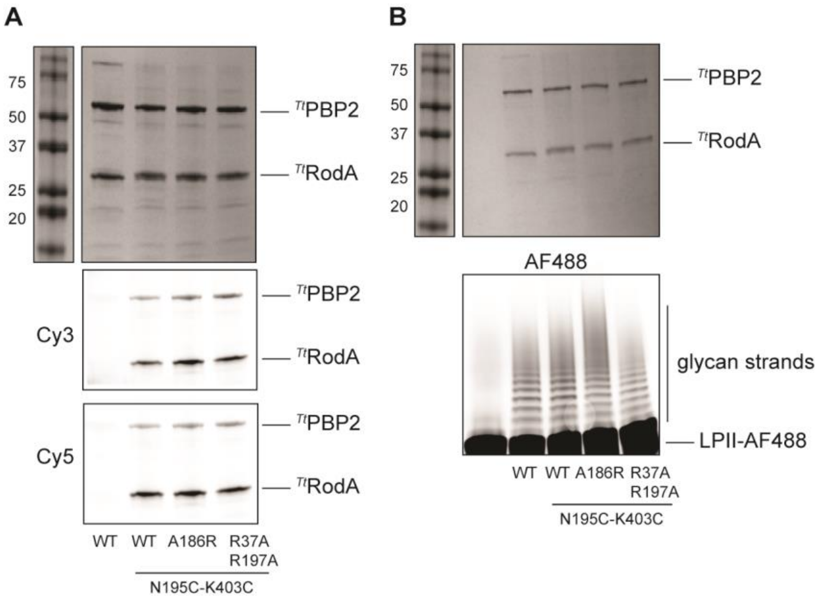


**Supplementary Figure 1.** Biochemical validation of $^{Tt}$RodA-PBP2 smFRET imaging constructs. A. Gels of fluorophore-labeled double-cysteine imaging mutants and mock-labeled $^{Tt}$WT control show specific incorporation of Cy3 and Cy5 fluorophores in RodA and PBP2 (pMS239, pSI7, pSI12, pSI13). Top: coomassie-stained control; middle: Cy3 signal; bottom: Cy5 signal. B. Imaging mutants retain activity after labeling (same constructs as A). Top: coomassie-stained control; bottom: Cy2 signal, showing polymerization of AF488-labeled lipid II.

**Supplementary Table 2. smFRET trajectory parameters**

| | Construct | Trajectories | Observations | Transitions | PDF fits (means) |
|---|---|---|---|---|---|
| $^{Tt}$RodA-PBP2 | WT | 222 | 15,984 | 124* | 0.83 |
| | R37A-R197A | 436 | 33,665 | 540 | 0.49<br>0.8 |
| | A186R | 415 | 38,978 | 779 | 0.45<br>0.75 |
| $^{Ec}$RodA-PBP2 | WT | 270 | 59,064 | 2045 | 0.52<br>0.78 |
| | ALFA | 94 | 17,423 | 400 | 0.60<br>0.87 |
| | ALFA + Nb-Fab | 122 | 30,341 | 4196 | 0.38<br>0.63<br>0.77 |
| | T52A | 173 | 66,054 | 2684 | 0.5<br>0.78 |
| | L61R | 100 | 35,745 | 1886 | 0.42<br>0.67<br>0.90 |
| | T52R-S54A-N55E | 195 | 45,262 | 1048 | 0.16<br>0.45 |

*Note that virtually none of $^{Tt}$WT "transitions" are bona fide, i.e., they are an artifact of fitting the data to a two-state model and correspond to exchange within the high-FRET state, rather than to true cross-state exchange.

**Supplementary Table 3. Cryo-EM sample preparation and data collection**

| | $^{Tt}$RodA-PBP2$^{WT}$ | $^{Tt}$RodA-PBP2$^{A186R}$ | $^{Ec}$RodA-PBP2$^{WT}$ |
|---|---|---|---|
| Grid type | C-flat holey carbon | C-flat holey carbon | UltrAuFoil holey gold |
| Protein concentration | 3.5 mg/mL | 3 mg/mL | 3 mg/mL |
| Blotting parameters | 7 s blot time<br>15 blot force | 7 s blot time<br>15 blot force | 7 s blot time<br>15 blot force |
| Microscope | Titan Krios | Titan Krios | Titan Krios |
| Pixel size (Å) | 0.825 | 0.825 | 0.825 |
| Voltage (kV) | 300 | 300 | 300 |
| Detector | K3 | K3 | K3 |
| Total exposure (e-/Å) | 64 | 58 | 59 |
| Defocus range (μm) | −1 to −2.5 | −1.2 to −2.5 | −1.2 to −2.5 |
| Number of frames per micrograph | 50 | 54 | 57 |
| Exposure time (s) | 3 | 1.4 | 2 |
| Final particles | 93,553 | 78,333 (open)<br>92,393 (closed) | 78,031 (1:1 protein: micelle dimer) |
| Symmetry | C1 | C1 | C1 |
| Nominal resolution (Å) | 6.5 | 6.1 (closed)<br>NA (open) | 9 |

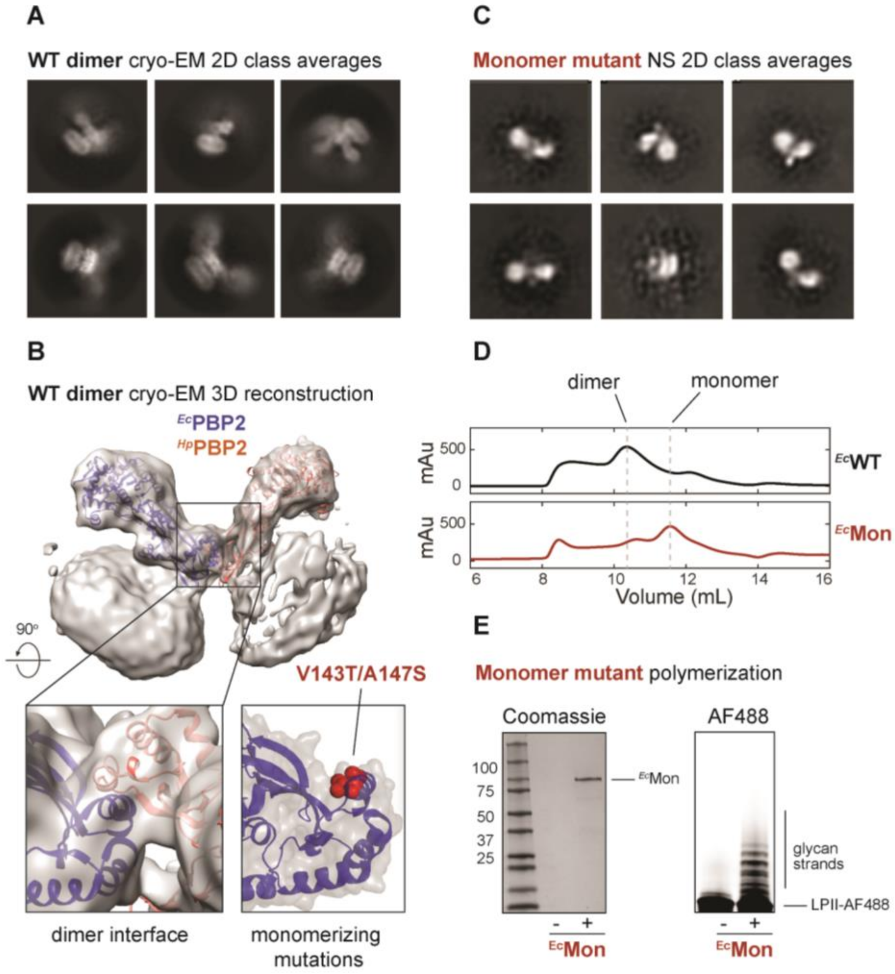


**Supplementary Figure 4.** Designing a monomeric $^{Ec}$RodA-PBP2 construct.
A. Representative cryo-EM 2D class averages of $^{Ec}$RodA-PBP2$^{WT}$ fusion construct (pSS50) show that the protein forms 2 types of dimers (1:1 protein:micelle and 2:1 protein:micelle). Each class contains 2-2.5k particles. B. 3D reconstruction of the main population of 1:1 protein:micelle dimeric particles. The conformation of the pedestal domain of PBP2 appeared distinct in the two protomers: either with the anchor and head domains closed (blue), similar to X-ray structure of ecto-$^{Ec}$PBP2 (PDB: 6g9f), or open (orange), similar to the MreC-bound state of ecto-$^{Hp}$PBP2 (PDB: 5lp5). To resolve the dimer interface, $^{Ec}$PBP2 and $^{Hp}$PBP2 crystal structures were fit into the cryo-EM map, assuming a symmetric dimer (bottom left). Mutations (red spheres) were designed to disrupt dimerization by increasing the hydrophilicity of the dimerization helix (right). C. Representative negative-stain electron microscopy 2D class averages show that $^{Ec}$RodA-PBP2$^{V143T\text{-}A147S}$ mutant is monomeric ($^{Ec}$Mon, pSI108). Each class contains ~500 particles. D. V143T-A147S mutations shift the size-exclusion elution profile of $^{Ec}$RodA-PBP2. E. Monomeric mutant retains polymerization activity. Left: coomassie-stained control; right: Cy2 excitation, showing polymerization of AF488-labeled lipid II.

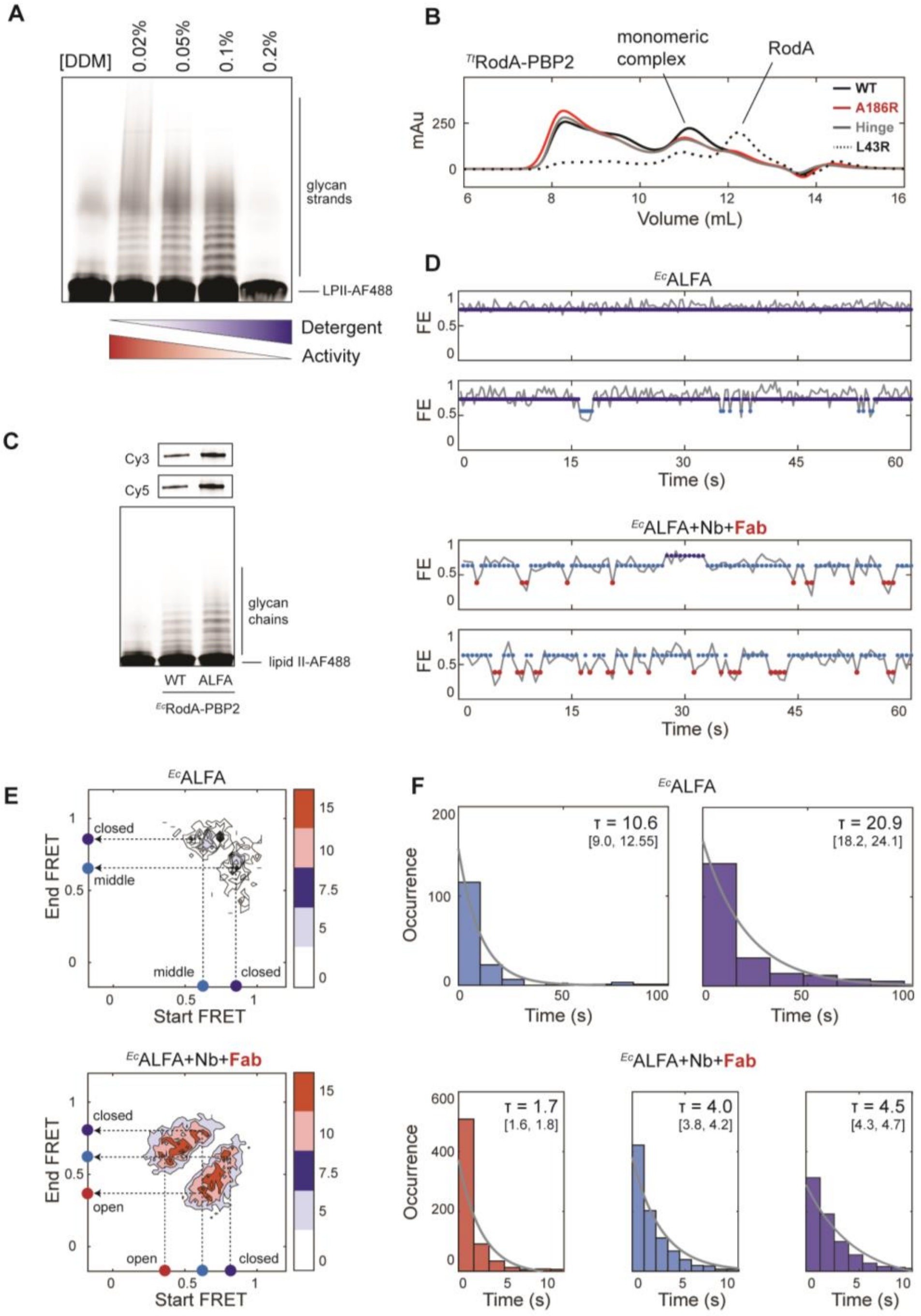


**Supplementary Figure 5.** Biochemical validation of constructs used in polymerization assays.
A. Detergent concentration dramatically affects polymerization activity. Polymerization activity of $^{Ec}$RodA-PBP2$^{Mon}$ (pSI126) assayed under a range of DDM concentrations above CMC (0.02-0.2%). AF488-labeled glycan chains were visualized with Cy2 excitation. B. Size-exclusion elution (SEC) profiles of *T. thermophilus* RodA-PBP2 WT and mutants (pSI7, pSI12, pSI13, pMS292). All PBP2 mutants retain WT fold and form a stable complex with RodA except for L43R, which shows considerable dissociation into $^{Tt}$RodA and $^{Tt}$PBP2. C. $^{Ec}$RodA-PBP2$^{ALFA}$ retains activity after labeling. Top: Cy3 and Cy5 signals, showing fluorophore labeling; bottom: Cy2 signal, showing polymerization of AF488-labeled lipid II. D. Representative trajectories of $^{Ec}$ALFA with 1:1 Nb:Fab complex from Fig. 4C. Mean values of states are marked. E. Transition density plots, normalized to total observation time, show that dynamic exchange into the low-FRET state is dramatically increased upon addition of the Fab-Nb complex. F. Dwell time histograms and exponential fits for the low-FRET (red), middle-FRET (light blue) and high-FRET (blue) states for $^{Ec}$ALFA with or without Nb-Fab. Mean dwell times alongside 95% confidence intervals for all populations are indicated on the plots.

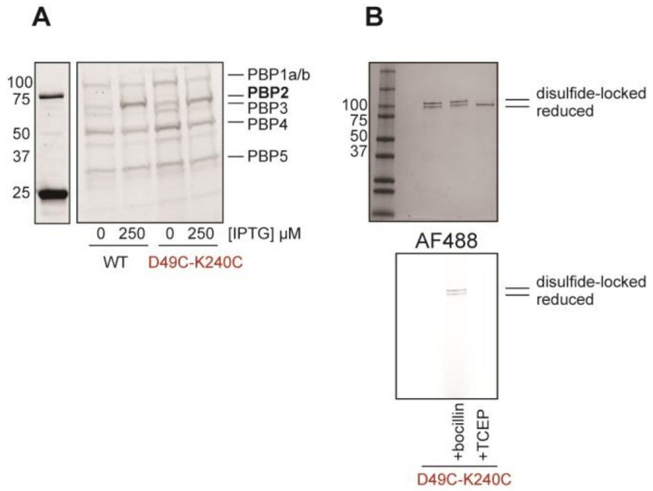


**Supplementary Figure 6.** Disulfide lock does not impair protein expression or fold.

A. ${}^{Ec}$RodA-PBP2$^{D49C\text{-}K240C}$ mutant expresses at 0.5-1x of ${}^{Ec}$RodA-PBP2$^{WT}$. Bocillin labeling was used to visualize and quantify PBP2 expression levels upon induction with IPTG in complementation strains FB38/pHC857 and FB38/pSI47 (as in Fig. 5C). Image shown is representative of 2 biological replicates. C. ${}^{Ec}$PBP2$^{D49C\text{-}K240C}$ (pSI128) forms a disulfide with ~50% efficiency. Top: SDS-PAGE shows that ${}^{Ec}$D49C-K240C sample contains a second band, in ${}^{Ec}$Mon (compare with Fig. S3E), which disappears upon treatment with TCEP. The top and bottom bands were assigned to the disulfide-locked and reduced species, respectively. The efficiency of disulfide formation was quantified from the relative intensities of the two bands. Bottom: bocillin labeling visualized with Cy2 excitation.

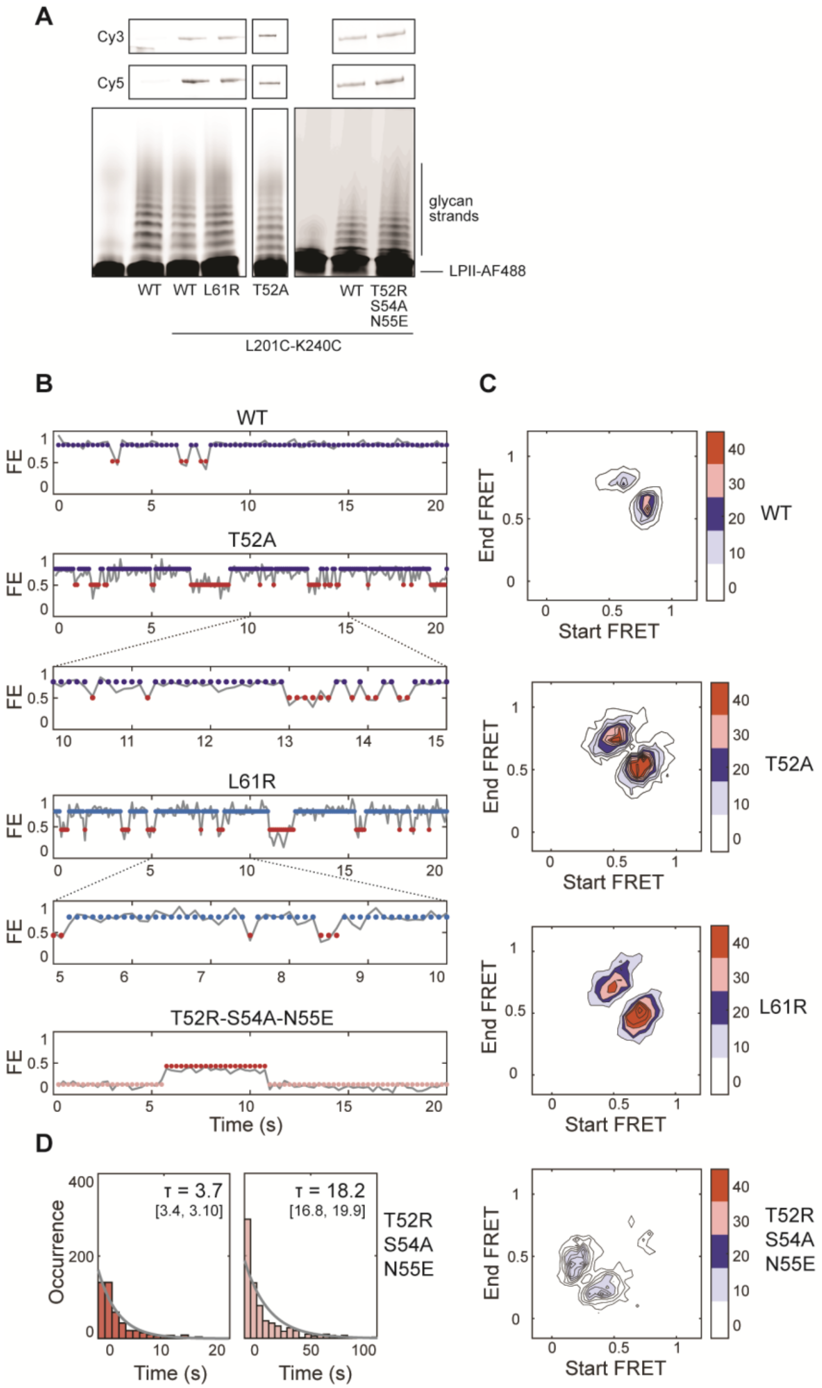


**Supplementary Figure 7.** Biochemical validation of $^{Ec}$RodA-PBP2 smFRET imaging constructs. A. smFRET imaging constructs are specifically labeled with Cy3 and Cy5 fluorophores and retain activity after labeling. Top: Labeling gels of the double-cysteine imaging mutants and the mock-labeled $^{Ec}$RodA-PBP2$^{WT}$ control (pSI131, pSI126, pSI127, pSI149, pSI152). Bottom: Lipid II-AF488 polymerization assay with the same constructs. B. Representative trajectories of smFRET constructs from Fig. 6B. $^{Ec}$RodA-PBP2$^{Mon}$ trajectory is reproduced from Fig. 3 for comparison. The insets show expansions of $^{Ec}$T52A and $^{Ec}$L61R trajectories to highlight individual transition events. Note that these mutants were imaged at a higher frame rate than the rest of the constructs (10 $s^{-1}$ vs 4 $s^{-1}$) to capture their faster dynamics. C. Transition plot analysis shows a comparison of transition frequency between dynamic mutants and $^{Ec}$Mon (reproduced from Fig. 3C), normalized to total observation time. D. Dwell time histograms and exponential fits for the lowest-FRET (pink), and low-FRET (red) states for $^{Ec}$RodA-PBP2$^{T52R\text{-}S54A\text{-}N55E}$. Mean dwell times alongside 95% confidence intervals for all populations are indicated on the plots.

**Supplementary Table 8: bacterial strains used for *in vivo* assays**

| **Strain** | **Genotype** | **Source** |
| --- | --- | --- |
| MG1655 | *rph1 ilvG rfb-50* | Guyer et al. 1981 |
| MG1655/pSI56 | *rph1 ilvG rfb-50* / $P_{tac}$:*pbpA-rodA* | This study |
| MG1655/pSI57 | *rph1 ilvG rfb-50* / $P_{tac}$:*pbpA(D49C-K240C)-rodA* | This study |
| MG1655/pSI58 | *rph1 ilvG rfb-50* / $P_{tac}$:*pbpA(D49C)-rodA* | This study |
| MG1655/pSI59 | *rph1 ilvG rfb-50* / $P_{tac}$:*pbpA(K240C)-rodA* | This study |
| MG1655/pSI63 | *rph1 ilvG rfb-50* / $P_{tac}$:*pbpA(S330A)-rodA* | This study |
| MG1655/pHC857 | *rph1 ilvG rfb-50* / $P_{lac}$:*pbpA-rodA* | This study |
| MG1655/pHC858 | *rph1 ilvG rfb-50* / $P_{lac}$:*pbpA(S330A)-rodA* | This study |
| MG1655/pSI47 | *rph1 ilvG rfb-50* / $P_{lac}$:*pbpA(D49C-K240C)-rodA* | This study |
| MG1655/pSI48 | *rph1 ilvG rfb-50* / $P_{lac}$:*pbpA(D49C)-rodA* | This study |
| MG1655/pSI49 | *ph1 ilvG rfb-50* / $P_{lac}$:*pbpA(K240C)-rodA* | This study |
| MG1655/pSI154 | *rph1 ilvG rfb-50* / $P_{lac}$:*pbpA(T52A)-rodA* | This study |
| MG1655/pSI157 | *rph1 ilvG rfb-50* / $P_{lac}$:*pbpA(T52R-S54A-N55E)-rodA* | This study |
| MG1655/pRY47 | *rph1 ilvG rfb-50* / $P_{lac}$: empty | This study |
| FB38 | *rph1 ilvG rfb-50 ΔlacIZYA::frt mrdAB::aph* | Bendezu & de Boer, 2008 |
| FB38/pHC857 | *rph1 ilvG rfb-50 ΔlacIZYA::frt mrdAB::aph* / $P_{lac}$:*pbpA-rodA* | P1(FB38) x MG1655/pHC857 |
| FB38/pHC858 | *rph1 ilvG rfb-50 ΔlacIZYA::frt mrdAB::aph* / $P_{lac}$:*pbpA(S330A)-rodA* | P1(FB38) x MG1655/pHC858 |
| FB38/pSI47 | *rph1 ilvG rfb-50 ΔlacIZYA::frt mrdAB::aph* / $P_{lac}$:*pbpA(D49C-K240C)-rodA* | P1(FB38) x MG1655/pSI47 |
| FB38/pSI48 | *rph1 ilvG rfb-50 ΔlacIZYA::frt mrdAB::aph* / $P_{lac}$:*pbpA(D49C)-rodA* | P1(FB38) x MG1655/pSI48 |
| FB38/pSI49 | *rph1 ilvG rfb-50 ΔlacIZYA::frt mrdAB::aph* / $P_{lac}$:*pbpA(K240C)-rodA* | P1(FB38) x MG1655/pSI49 |
| PR5 | *rph1 ilvG rfb-50 mreC(R292H) yrdE-kan* | Rohs et al. 2018 |

| | | |
|---|---|---|
| PR5/pHC857 | *rph1 ilvG rfb-50 mreC(R292H) yrdE-kan* / $P_{lac}$:*pbpA-rodA* | P1(PR5) x MG1655/pHC857 |
| PR5/pSI154 | *rph1 ilvG rfb-50 mreC(R292H) yrdE-kan* / $P_{lac}$:*pbpA(T52A)-rodA* | P1(PR5) x MG1655/pSI154 |
| PR5/pSI157 | *rph1 ilvG rfb-50 mreC(R292H) yrdE-kan* / $P_{lac}$:*pbpA(T52R-S54A-N55E)-rodA* | P1(PR5) x MG1655/pSI157 |
| PR5/pRY47 | *rph1 ilvG rfb-50 mreC(R292H) yrdE-kan* / $P_{lac}$:*empty* | P1(PR5) x MG1655/pRY47 |

**Supplementary Table 9: plasmids used for *in vivo* assays**

| **Plasmid** | **Genotype** | **ori** | **Source** |
|---|---|---|---|
| pHC857 | *cat* $lacI^q$ $P_{lac}$:*pbpA-rodA* | pBR/colE1 | Cho et al. 2014 |
| pHC858 | *cat* $lacI^q$ $P_{lac}$:*pbpA(S330A)-rodA* | pBR/colE1 | Cho et al. 2014 |
| pSI47 | *cat* $lacI^q$ $P_{lac}$:*pbpA(D49C-K240C)-rodA* | pBR/colE1 | This study |
| pSI48 | *cat* $lacI^q$ $P_{lac}$:*pbpA(D49C)-rodA* | pBR/colE1 | This study |
| pSI49 | *cat* $lacI^q$ $P_{lac}$:*pbpA(K240C)-rodA* | pBR/colE1 | This study |
| pHC800 | *cat* $lacI^q$ $P_{tac}$ | pBR/colE1 | Cho et al. 2014 |
| pSI56 | *cat* $lacI^q$ $P_{tac}$:*pbpA-rodA* | pBR/colE1 | This study |
| pSI57 | *cat* $lacI^q$ $P_{tac}$:*pbpA(D49C-K240C)-rodA* | pBR/colE1 | This study |
| pSI58 | *cat* $lacI^q$ $P_{tac}$:*pbpA(D49C)-rodA* | pBR/colE1 | This study |
| pSI59 | *cat* $lacI^q$ $P_{tac}$:*pbpA(K240C)-rodA* | pBR/colE1 | This study |
| pSI63 | *cat* $lacI^q$ $P_{tac}$:*pbpA(S330A)-rodA* | pBR/colE1 | This study |
| pSI154 | *cat* $lacI^q$ $P_{lac}$:*pbpA(T52A)-rodA* | pBR/colE1 | This study |
| pSI157 | *cat* $lacI^q$ $P_{lac}$:*pbpA(T52R-S54A-N55E)-rodA* | pBR/colE1 | This study |
| pRY47 | *cat* $lacI^q$ $P_{lac}$: empty | pBR/colE1 | Cho et al. 2014 |

**Supplementary Table 10: plasmids used for *in vitro* assays**

| Plasmid | Genotype | Source |
|---|---|---|
| pMS239 | *ColA-T7:TtPBP2-3C-PrtC; $P_{T7}$:His6-SUMO-Flag-3C-TtRodA* | Sjodt et al. 2020 |
| pMS292 | *ColA-T7:TtPBP2(L43R)-3C-PrtC; $P_{T7}$:His6-SUMO-Flag-3C-TtRodA* | |
| pMS293 | *ColA-T7:TtPBP2(A186R)-3C-PrtC; $P_{T7}$:His6-SUMO-Flag-3C-TtRodA* | Sjodt et al. 2020 |
| pSI7 | *ColA-T7:TtPBP2(K403C)-3C-PrtC; $P_{T7}$:His6-SUMO-Flag-Avitag-TtRodA(N195C)* | This study |
| pSI12 | *ColA-T7:TtPBP2(A186R-K403C)-3C-PrtC; $P_{T7}$:His6-SUMO-Flag-Avitag-TtRodA(N195C)* | This study |
| pSI13 | *ColA-T7:TtPBP2(R37A-R197A-K403C)-3C-PrtC; $P_{T7}$:His6-SUMO-Flag-Avitag-TtRodA(N195C)* | This study |
| pSS50 | *pAM205-T7:His6-SUMO-Flag-3C-EcPBP2-GGGSx3-EcRodA* | Rohs et al. 2018 |
| pSI108 | *T7:His6-SUMO-Flag-3C-EcPBP2(V143T-A147S)-GGGSx3-EcRodA* | This study |
| pSI126 | *T7:His6-SUMO-Flag-3C-EcPBP2(V143T-A147S-R425C)-GGGSx3-EcRodA(C133A-L201C)* | This study |
| pSI147 | *T7:His6-SUMO-Flag-3C-EcPBP2(V143T-A147S-[I182-ALFA tag-A201] R425C)-GGGSx3-EcRodA(C133A-L201C)* | This study |
| pSI127 | *T7:His6-SUMO-Flag-3C-EcPBP2(L61R-V143T-A147S-R425C)-GGGSx3-EcRodA(C133A-L201C)* | This study |
| pSI128 | *T7:His6-SUMO-Flag-3C-EcPBP2(D49C-K240C-V143T-A147S)-GGGSx3-EcRodA(C133A)* | This study |
| pSI129 | *T7:His6-SUMO-Flag-3C-EcPBP2(D49C-V143T-A147S)-GGGSx3-EcRodA(C133A)* | This study |
| pSI130 | *T7:His6-SUMO-Flag-3C-EcPBP2(K240C-V143T-A147S)-GGGSx3-EcRodA(C133A)* | This study |
| pSI131 | *T7:His6-SUMO-Flag-3C-EcPBP2(V143T-A147S)-GGGSx3-EcRodA(C133A)* | This study |
| pSI149 | *T7:His6-SUMO-Flag-3C-EcPBP2(T52A-V143T-A147S-R425C)-GGGSx3-EcRodA(C133A-L201C)* | This study |

| | | |
|---|---|---|
| pSI152 | *T7:His6-SUMO-Flag-3C-EcPBP2(T52R-S54A-N55E-V143T-A147S-R425C)-GGGSx3-EcRodA(C133A-L201C)* | This study |
| pSI145 | *pET26b-T7:PelB-NbALFA-6xHis* | This study, nanobody sequence from Götzke & Kilisch et al. 2019 |
| pSI146 | *pET26b-T7:PelB-NbALFA(Q116K-Q119P)-6xHis* | This study |
| pMAS478 | *pTarget:* heavy chain | This study, Fab sequence from Bloch et al. 2021 |
| pMAS481 | *pD2610-v5:* light chain | This study, Fab sequence from Bloch et al. 2021 |
